# Aggregated *Mycobacterium tuberculosis* enhances the inflammatory response

**DOI:** 10.1101/2021.03.23.436577

**Authors:** Hylton E. Rodel, Isabella Markham Ferreira, Carly G.K Ziegler, Yashica Ganga, Mallory Bernstein, Shi-Hsia Hwa, Kievershen Nargan, Gila Lustig, Gilla Kaplan, Mahdad Noursadeghi, Alex K. Shalek, Adrie Steyn, Alex Sigal

## Abstract

*Mycobacterium tuberculosis* (Mtb) readily aggregates in culture and Mtb aggregates in the lung were observed in experimental Mtb infection. However, the physiological consequences of Mtb aggregation are incompletely understood. Here we examined the human macrophage transcriptional response to aggregated Mtb relative to infection with non-aggregated single or multiple bacilli per host cell. Infection with aggregated Mtb led to an early upregulation of pro-inflammatory associated genes and enhanced TNF*α* signaling via the NF*κ*B pathway. Both these pathways were significantly upregulated relative to infection with single bacilli, and TNF*α* signaling was also significantly elevated relative to infection with multiple non-aggregated Mtb. Secretion of TNF*α* and downstream cytokines were also enhanced. On a longer timescale, aggregate infection led to overall increased acidification per macrophage and a high proportion of death in these cells after aggregate phagocytosis. Host cell death did not occur when Mtb aggregates were heat killed despite such clumps being readily picked up. To validate that Mtb aggregates do occur in the human lung, we document Mtb aggregates surrounding a cavity in a human TB lesion. Aggregates may therefore be present in some lesions and elicit a stronger inflammatory response resulting in recruitment of additional phagocytes and their subsequent death, potentially leading to necrosis and transmission.

## Introduction

Mtb infection of a human lung results in either a latent disease state, in which an individual is infected and asymptomatic, or active TB disease, manifesting as hemoptysis, lung damage, weight loss, and other severe systemic effects [1, 2, 3, 4, 5]. In active TB, Mtb infection leads to necrosis of the granuloma, the structure which encapsulates Mtb infection, attempting to isolate it from the surrounding lung [6, 4, 7, 8]. This in turn results in the necrotic granuloma entering the airway and expectoration of bacilli out of the lung and transmission to other hosts [1, 2, 3, 4].

The host-pathogen interactions that tip the balance to active disease are not clearly defined. Aggregation has been proposed to be associated with pathogenicity [9, 10] and increase Mtb virulence in mice and *ex vivo* [11, 12, 13]. Mtb aggregates have been observed in human lung tissue [7], though the fraction of bacilli in this state is unclear. Additionally, Mtb aggregates have recently been shown to be transmitted in bio-aerosols [14].

We have previously demonstrated using time-lapse microscopy that infection with aggregated Mtb preferentially leads to macrophage death [15] and this has been observed by others using different methods [16, 17]. Death of infected macrophages in turn results in rapid replication of the bacilli inside the dead infected cells [15], an observation which was independently confirmed [18]. The necrotic, infected cell may then be phagocytosed by another macrophage, leading to additional cycles of host cell death and bacterial growth which may close a positive-feedback loop [15], provided more phagocytes are recruited to the infection area.

Macrophages show extensive transcriptional remodeling of their immune and inflammatory pathways [19, 20, 21]. This includes upregulation and secretion of TNF*α*, IL8, CCL3, CCL4, IL1*β* and other factors involved in inflammation [22, 23, 24]. This response activates macrophages for the killing of phagocytosed Mtb [25] and recruits additional cell types, such as neutrophils, to the site of Mtb infection [26, 27]. While such a response may be host protective, macrophage cell death may also be a consequence[25, 28, 9]. Here we investigated the effects of Mtb aggregation on the macrophage early transcriptional response. We observed that infection with Mtb aggregates led to a stronger early inflammatory response in human monocyte derived macrophages, with higher secretion of TNF*α*, as well as upregulation of genes leading to chemotaxis. We also observed that Mtb aggregates accounted for a substantial number of the Mtb identified on the periphery of a cavitary lesion. Taken together, these results may be consistent with Mtb aggregation playing a role in TB pathogenesis.

## Results

### Aggregated Mtb shows a distinct transcriptional response dependent on both Mtb number and aggregation state

We asked whether bacterial aggregation could modulate the early host cell transcriptional response, where secondary processes associated with host cell death have not yet begun.

We infected monocyte derived macrophages (MDM) with single or aggregated bacilli (3 repeats for each of 5 donors, *n* = 15). Single Mtb were obtained by bead beating Mtb grown in the absence of detergent, followed by filtration (Materials and methods). At 3 hours post-infection, we sorted live MDM which internalized aggregated Mtb (Figure 1A), or live MDM which internalized single Mtb or multiple single Mtb. As a control, we sorted uninfected MDM. We performed population RNA-Seq on the different sorted populations. We identified the most variably expressed genes after normalization and batch correction and used principal component analyses (PCA) to plot the top 0.1 % of variable genes across the different conditions. We observed that the infection conditions could be separated along the first principal component, where there was a graded response in gene expression from uninfected to aggregate infected cells (Figure 1B), indicating that both the number of infecting bacilli and the aggregation state determines the transcriptional profile. The genes contributing to this separation included TNF, IL8, CCL4, IL1*β*, CXCL2, CXCL3 and other genes involved in inflammation and chemotaxis (Figure 1C).

**Figure 1:**
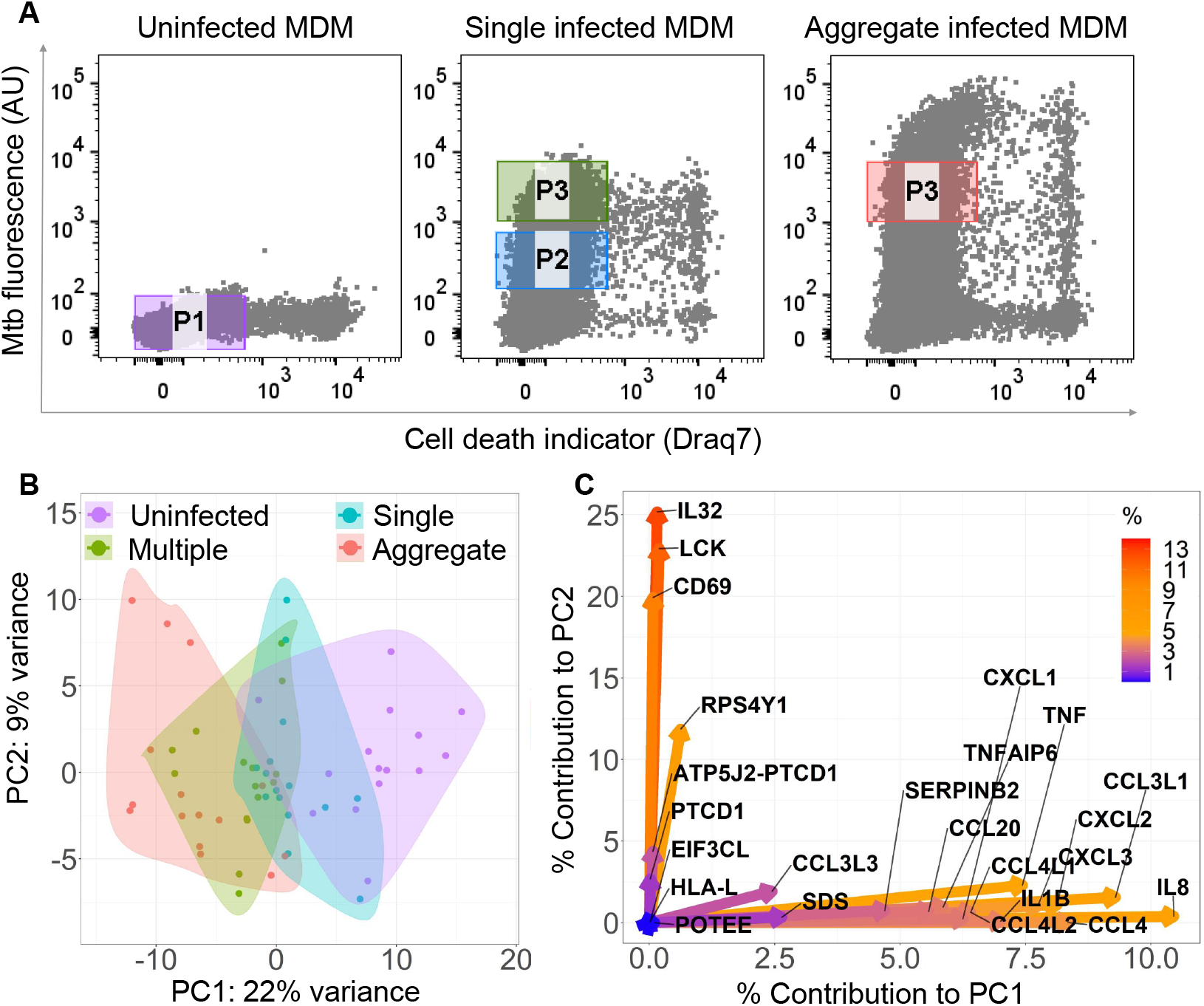
Mtb Aggregation changes the macrophage transcriptional response. (A) MDM were infected with mCherry labeled, aggregated Mtb (right panel), single Mtb bacilli (middle panel) or uninfected (left panel) and sorted for RNA-Seq 3 hours post-infection. Populations consisted of cells infected with aggregated Mtb (Gate P3 -right panel), cells infected with single or few bacilli (Gate P2 -middle panel), cells infected with multiple single bacilli (Gate P3 -middle panel), or uninfected (Gate P1 -left panel). Dead cells were excluded. X-axis is signal from the cell death detection dye DRAQ7, y-axis is mCherry signal from Mtb infection. (B) Principal component analysis (PCA) of the top 0.1% most variably expressed genes following rlog normalization and batch correction in R. Small circles are individual experiments (3 repeats from each of 5 blood donors). (C) Percentage contribution of individual genes used in the PCA. Color bar represents the percent contribution of individual genes to the first two principal components.

We examined the functional differences between infection conditions at the transcriptional level using the Gene Set Enrichment Analysis (GSEA). We compared all infection conditions relative to each other and plotted the normalized enrichment scores (NES) for two gene sets (“TNF*α* signaling via NF-*κ*B” and “Inflammatory response”) that were most significantly different between comparisons (Figure 2A-B, Table S1). We observed that, relative to uninfected cells, there was enrichment of both gene sets when MDM were infected with single Mtb. However, a further increase in the enrichment score occurred when infection was by multiple non-aggregated bacilli and was highest with aggregated bacilli. Interestingly, both the number of bacilli and aggregation state showed an effect: there was enrichment in both TNF*α* signaling and the inflammatory response with multiple non-aggregated Mtb relative to single Mtb, and in TNF*α* signaling with aggregate infection relative to infection with multiple non-aggregated bacilli per cell (Figure 2A).

**Figure 2:**
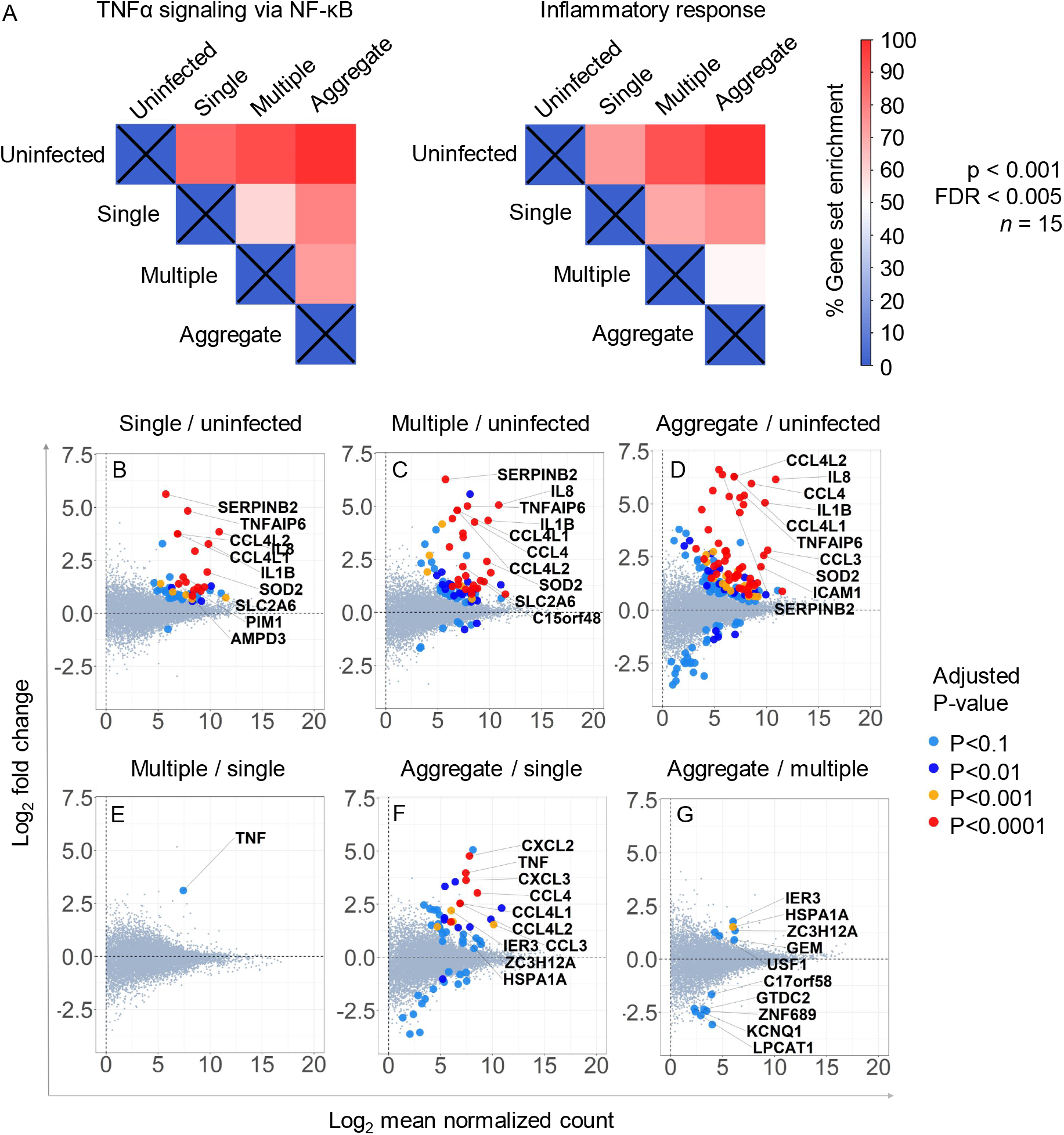
Gene sets and individual genes differentially regulated between infection conditions. Normalised enrichment score (NES), expressed as percentage of maximum enrichment for the gene sets defined as “TNF*α* signaling via NF-*κ*B”, and “Inflammatory response”. Enrichment scores were calculated for all treatment comparisons and were significantly different at Nominal p *<* 0.001 and FDR *<* 0.005, with the exception of the aggregate to multiple comparison for the “inflammatory response” gene set (where p *<* 0.05 and FDR = 0.24). Individual differentially regulated genes were identified by comparing infection conditions (B-G) in DESeq2. Large circles represent genes differentially regulated between two infection conditions at a significance indicated in the corresponding colour key, while smaller grey circles are not significantly different. Fold-change in expression is on the y-axis, with values above zero indicating up-regulation and below zero down-regulation. Read count is on the x-axis.

We next investigated the specific genes which were significantly up or down-regulated under the different infection conditions. When MDM were infected with single Mtb, the transcriptional response predominantly consisted of an upregulation of a subset of genes, including the TNF*α* responsive genes IL1*β*, CCL4L1 and CCL4L2, SERPINB2, and TNFAIP6 (Figure 2B, Table S2). More genes were upregulated when multiple single Mtb bacilli infected one MDM, including additional TNF*α* responsive genes IL8 and CCL4 (Figure 2C). With aggregate infection, there was a highly upregulated cluster of genes, including CCL4L1 and CCL4L2, IL8, CCL4, IL1*β*, SERPINB2, and TNFAIP6 (Figure 2D). These genes had functions including inflammation, neutrophil chemotaxis and regulation of apoptosis [29, 30, 31, 32, 33].

A comparison of multiple to single infected MDM yielded enhancement in TNF*α* expression (Figure 2E). In contrast, comparison of aggregate to single infection included enhancement of TNF*α* and also CCL4, CCL4L1, CCL4L2, CXCL2, and CXCL3 (Figure 2F). Genes were both up and down regulated in the comparison between aggregate and multiple single infection (Figure 2G). The genes HSPA1A and IER3 have been shown to be involved in the negative regulation of apoptotic cell death [34, 35]. Taken together, there is a trend toward stronger expression of inflammatory mediators with a progression from single infection to infection with unaggregated multiple Mtb, to aggregates.

We examined whether the observed transcriptional regulation is also reflected in cytokine secretion using TNF*α* and two cardinal TNF*α* responsive genes. Both at the transcriptional level (Figure 3A) and at the level of cytokine secretion (Figure 3B), TNF*α* response showed higher expression and secretion in MDM infected by aggregated Mtb relative to single Mtb. Interestingly, IL8, and IL6 did not show a significant upgregulation or secretion when infection was by aggregated relative to single Mtb.

**Figure 3:**
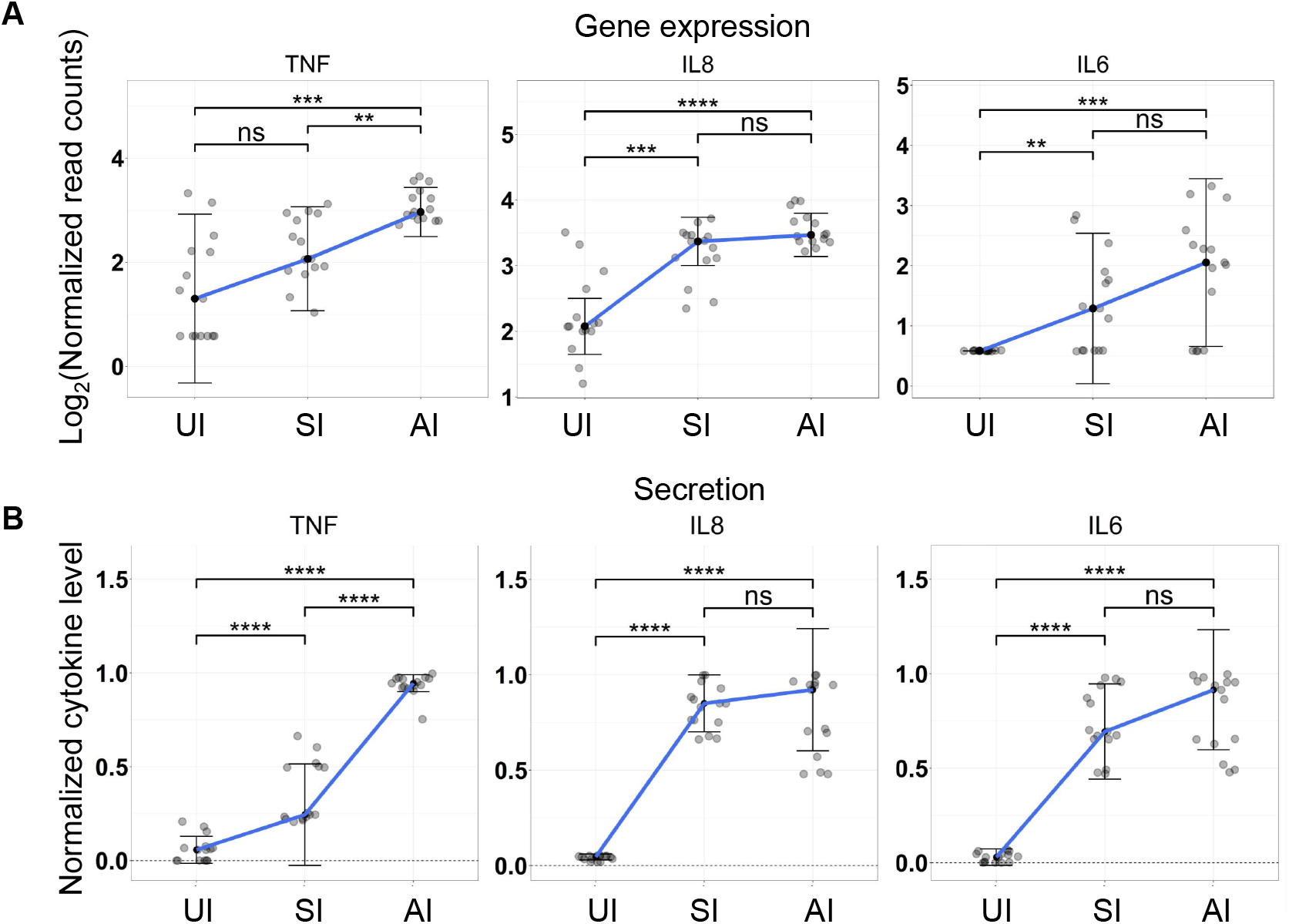
Transcriptional upregulation and secretion of TNF*α* and downstream genes with aggregated versus single Mtb infection. (A) Normalized transcripts or (B) cytokine secretion 3 hours post-Mtb infection. MDM were either uninfected (UI), infected with single Mtb (SI) or with aggregated Mtb (AI). Shown are median and IQR of the transcriptional or cytokine response from 15 independent infections of MDM from 5 blood donors. p-values are ** *<*0.01; *** *<*0.001; *** *<*0.0001; as determined by Mann-Whitney U test with Bonferroni multiple comparison correction.

### Mtb aggregates at the periphery of a TB cavity in the human lung

We analysed stained sections of lung tissue from a TB infected individual requiring clinically indicated lung lobe resection to determine if Mtb aggregates were present. Image analysis was performed using a custom image analysis code in Matlab 2019a (Materials and methods) to automatically identify Mtb within the tissue section. Mtb was classified as single or aggregated based on size and tested for cell association by proximity to adjacent cell nuclei (Figure S1). Mtb were located around the TB cavity but not elsewhere in the section (Figure 4A). A total of 1420 Mtb objects, containing different numbers of individual bacilli, were detected (Figure 4B). The total number of Mtb bacilli present in all objects was estimated at 2086, based on the average size of an individual bacterium (Materials and methods). 151 objects were classified as Mtb aggregates and corresponded to a minimum of 2.4 Mtb bacilli. These accounted for 28% of all detected Mtb bacilli (Figure 4C). 993 Mtb objects, 68% of all bacilli, detected were within close proximity of host cell nuclei and were classified as cell associated. 61% of the aggregated and 70% of the single Mtb bacilli were cell associated (Figure 4C). This data supports the existence of aggregates in a naturally infected human lung.

**Figure 4:**
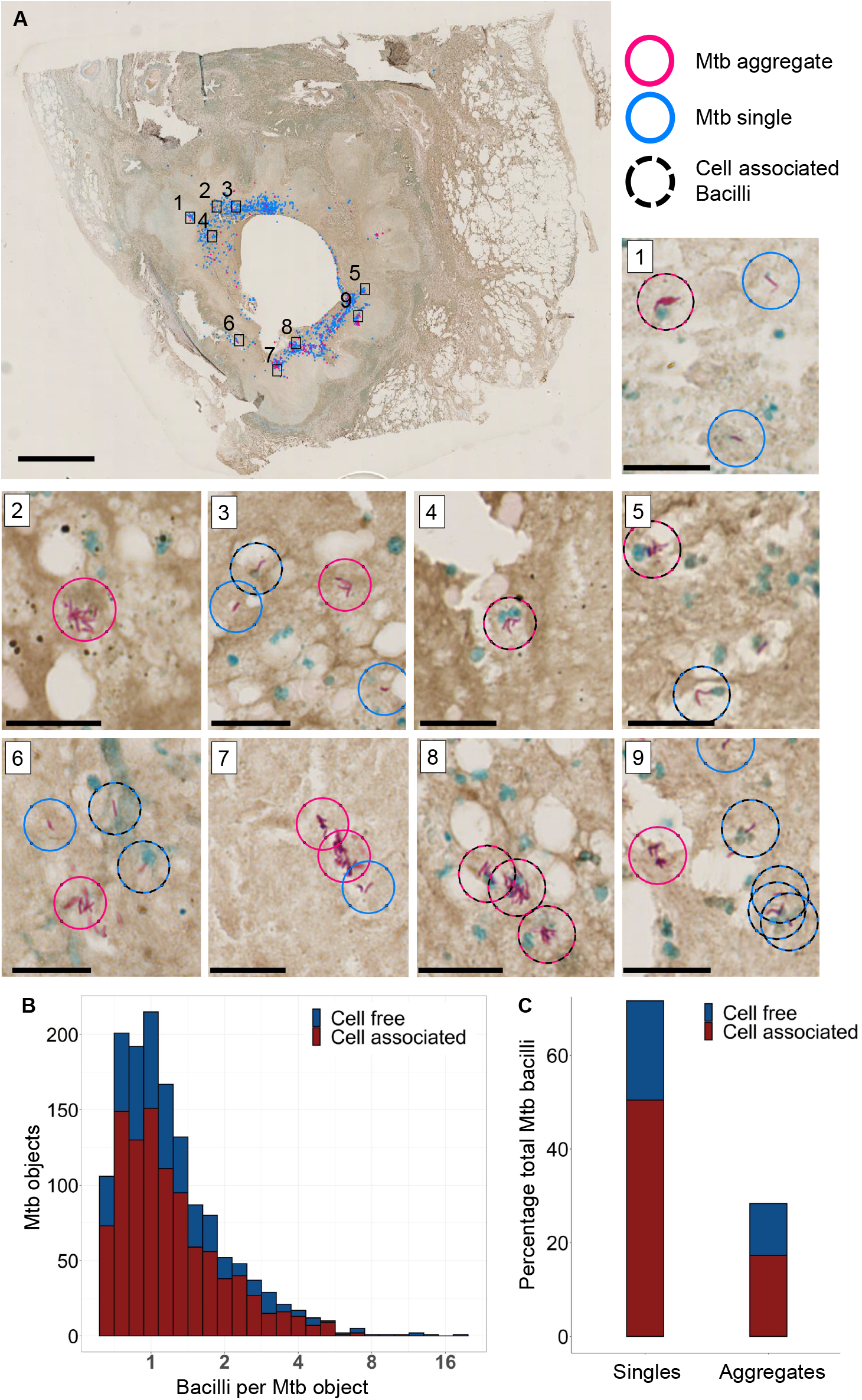
Mtb and Mtb aggregates are found near the periphery of a TB cavitary lesion. (A) Ziehl–Neelsen stain of a lung section. Aggregated bacilli are highlighted with a red circle and single bacilli with a blue circle. Scale bar is 3mm. A black dashed circle is overlaid if the Mtb are in close proximity to a cell nucleus (blue stain). Sub-areas 1-9 are magnified in separate panels. Scale bars are 20 *µ*m in the sub-areas. (B) Stacked histogram of the number of Mtb objects with varying numbers of Mtb bacilli found to be cell free or cell-associated (C) Stacked histogram of the total number of Mtb observed to be single bacilli or aggregates, and were found to be in close association with host cell nuclei.

### Aggregate mediated macrophage death requires live Mtb

We tested whether macrophage cell death, elicited by Mtb aggregates, required the bacilli to be live. MDM death was quantified using confocal fluorescence timelapse microscopy to detect penetration of the cell membrane permeability dye DRAQ7, where penetration of dye is associated with the loss of plasma membrane integrity (Figure 5A).

**Figure 5:**
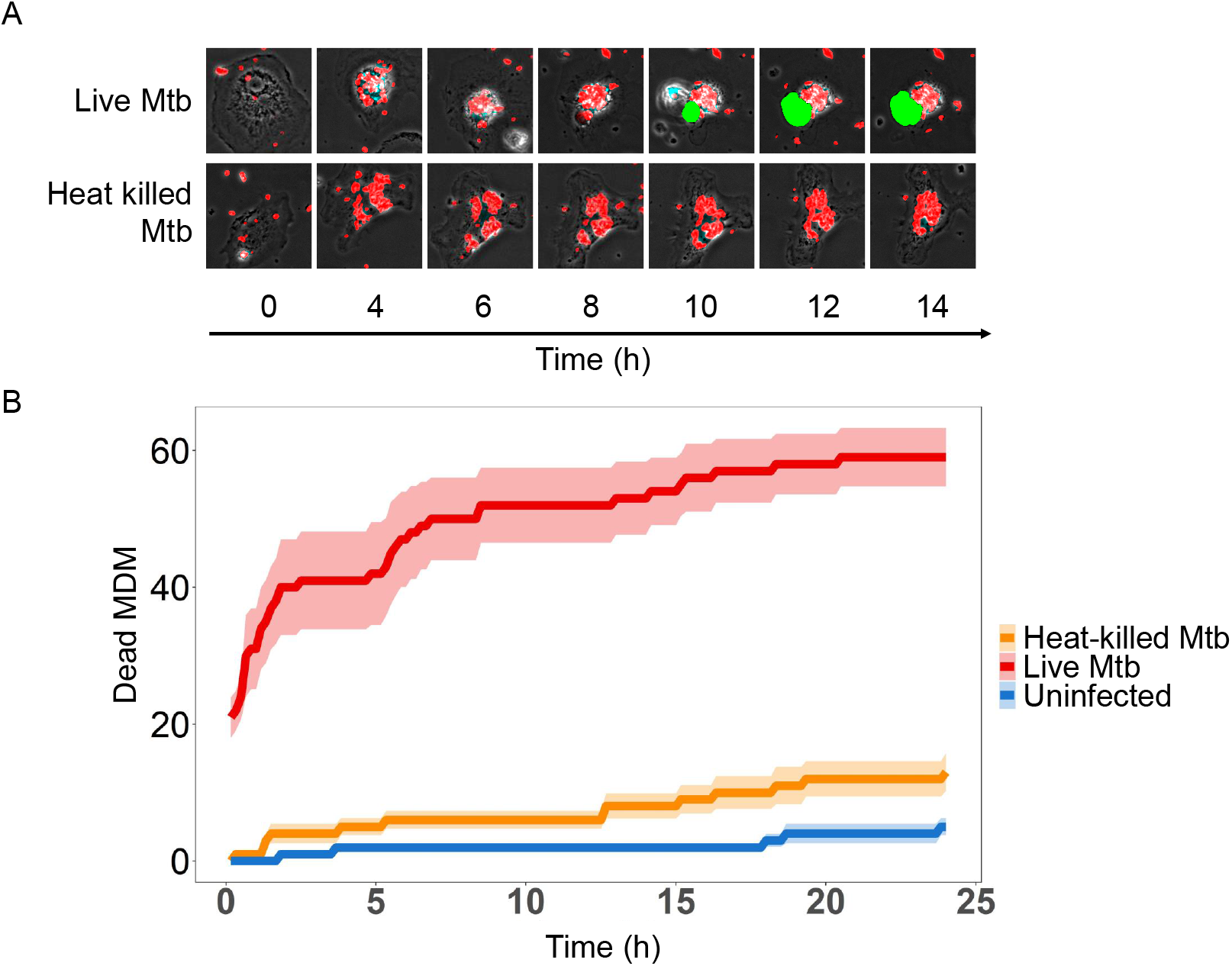
Macrophage death is dependent on infection with live Mtb aggregates. (A) Timelapse microscopy showing mCherry labelled Mtb (red) induced MDM death as detected by DRAQ7 (green). (B) The number of dead cells in MDM infected with live Mtb (red line), heat killed Mtb (orange), or uninfected (blue). Shown are mean *±*std of DRAQ7 positive cells per field of view at each timepoint measured.

We infected MDM with live Mtb aggregates (Figure 5A, Supplementary Movie 1) or with Mtb aggregates that had been heat-killed for 20 minutes at 80°C (Figure 5A, Supplementary Movie 2). We then monitored cell death in infected MDM over time. Despite the aggregates being heat killed, they were readily phagocytosed by macrophages (Supplementary Movie 2). We observed extensive MDM death when MDM were infected with live Mtb aggregates. In contrast, the number of dead MDM infected with fluorescent, heat killed aggregates did not markedly differ from uninfected MDMs (Figure 5B).

### Reduced acidification of bacilli in intracellular Mtb aggregates

To examine the possible causes of sub-optimal control of intracellular Mtb aggregates, which may lead to macrophage death, we asked whether the aggregation state modulates the ability of the macrophages to acidify phagosome associated Mtb. We infected MDM with single or aggregated Mtb for 6 hours, and imaged cells following staining with the LysoTracker reporter for acidification. We quantified Mtb and lysotracker fluorescence within each cell and the areas of overlap between Mtb and Lysotracker fluorescence (Figure 6A).

**Figure 6:**
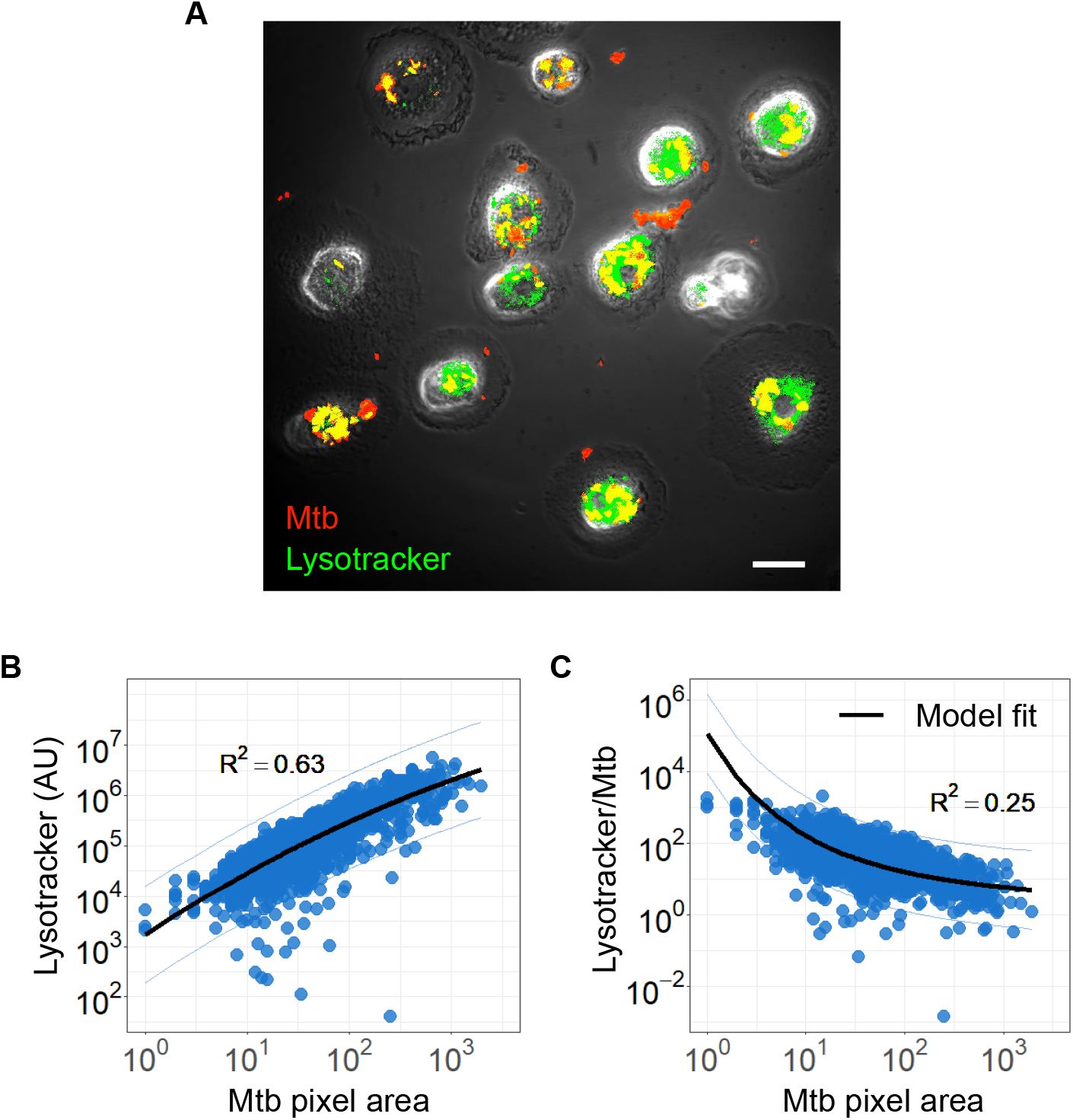
Mtb acidification per Mtb bacillus decreases with increasing Mtb aggregate size. (A) Image of lysotracker (green) colocalization with phagocytosed mCherry expressing Mtb (red). Scale bar is 20 *µ*m (B) Lysotracker fluorescence as a function of total aggregate area. The linear regression line is shown in black (*R*^2^ = 0.63, p*<*0.0001). (C) Ratio of Lysotracker fluorescence to Mtb fluorescence as a function of Mtb area. Black line shows a model based on the surface area to volume ratio (*R*^2^ = 0.25, p*<*0.0001, black line).

We plotted Lysotracker fluorescence against Mtb area and found that Lysotracker signal increases with Mtb signal (Figure 6B). However, the lysotracker signal normalised to Mtb fluorescence decreased as Mtb aggregate area increased (Figure 6C). Given that the phagosome closely envelopes the phagocytosed Mtb [36, 37], we hypothesized that the phagosome activity would be dependent on the surface area of the aggregate. The number of bacilli however, would be related to the phagosome volume, which would increase faster than its surface area as phagosome size increased. In support of this, fitting a surface area to volume ratio gave a reasonable fit of the data (*R*^2^ = 0.25, p*<*0.0001, black line in Figure 6C).

## Discussion

We investigated the early transcriptional response to macrophage infection by single or aggregated Mtb and observed that aggregate infection upregulates the TNF*α* and inflammatory responses relative to infection by single Mtb. This effect is mediated both by the number of bacilli infecting one cell, and the aggregation state, indicating that aggregation state in addition to the number of infecting bacilli per host cell drives the stronger response. The negative regulation of apoptosis in aggregate infected macrophages, relative to multiple infected macrophages, is consistent with our previous results indicating that apoptosis was not the cell death pathway elicited in macrophages infected by Mtb aggregates[15]. Interestingly, while TNF*α* itself is more strongly expressed and secreted at higher levels in aggregate Mtb infection, IL6 and IL8, key TNF*α* downstream genes are induced and secreted at similar levels independently of whether infection is by single or aggregated Mtb.

As we reported previously [15], we also observed here that aggregate infection tended to result in macrophage death. However, this occurred with live but not heat killed Mtb. The mechanism remains unclear, but may be consistent with an active process such as secretion of a toxin [38, 39].Possibly related to poorer control of aggregates and macropgage death, we observed that aggregates elicit stronger acidification of the phagosome. Yet, per Mtb bacillus in the aggregate, the acidification is reduced relative to that of phagosomes containing single bacilli, consistent with a decrease in the phagosome surface area to volume ratio. We speculate that while overall increased activation may be deleterious to the host cell, it has less cytopathic effect per bacillus in an aggregate.

One possibility of how Mtb aggregates may play a role in TB pathogenesis may be the recruitment of additional phagocyes to the infection site. CCL4 [40, 41], CXCL2 [41, 42, 43], and CXCL3 [44], upregulated in aggregate versus single infection (Figure 2H), are chemokines which recruit neutrophils. Neutrophil accumulation in Mtb infection leads to inflammation and has been shown to be detrimental to the host [27, 45]. Aggregated Mtb infection led to neutrophil accumulation in a mouse model [11]. Death of Mtb infected neutrophils may also enhance Mtb growth, which we have observed to be accelerated in dead infected cells [15] and may make aggregated Mtb a more virulent form of infection [12, 11, 46, 17, 10]. In support of this, we observed that Mtb aggregates occurred at a substantial frequency around a TB cavitary lesion. The aggregates would therefore be in the correct location for transmission. Mtb aggregates were recently observed in bio-aerosols [14]. If indeed aggregate transmission occurs, the increased virulence of Mtb aggregates may potentially increase the probability of developing active TB disease.

## Material and methods

### Ethical statement

Blood was obtained from adult healthy volunteers after written informed consent (University of KwaZulu-Natal Institutional Review Board approval BE022/13). Lung sections were obtained from clinically indicated resection due to TB complications (University of KwaZulu-Natal Institutional Review Board approval BE019/13). Written informed consent was obtained.

### Combination staining of human lungs

Human lung tissue were cut into 2 mm thick sections and picked on charged slides. Slides were baked at 56°C for 15 min. Mounted sections were dewaxed in xylene followed by rinse in 100% ethanol and 1 change of SVR (95%). Slides were then washed under running water for 2 min followed by antigen retrieval via heat induced epitope retrieval (HIER) in Tris-sodium chloride (pH 6.0) for 30 min. Slides were then cooled for 15 min and rinsed under running water for 2 min. Endogenous peroxide activity was blocked using 3% hydrogen peroxide for 10 min at room temperature (RT). Slides were then washed in PBST and blocked with protein block (Novolink) for 5 min at RT. Sections were incubated with primary antibodies for CD68 (M0814-CD68-KP1,DAKO,1:3000) followed by washing and incubation with the polymer (Novolink) for 30 min at RT Slides were then washed and stained with DAB for 5 min, washed under running water for 5min. For the combination staining, slides were incubated with heated carbol fuchsin for 10min and then washed in running tap water. 3% acid alcohol was applied to the slide to decolourize for 30 seconds or until sections appeared clear. Slides were then washed in running tap water for 2minutes and where then counter stained with methylene blue. Slides were rinsed under running water, dehydrated and mounted in Distyrene Plasticiser Xylene (DPX).

### Semi-automated detection of Mtb in histology slides

RGB Images of resected lung tissue in .ndpi format were converted to .TIFF file types (without compression) to enable compatibility with Matlab (Mathworks, Massachusetts, US) image processing functions. The resultant image files were sectioned into smaller tiles and imported individually into Matlab in a looped process. Saturated image tiles (tiles that had near completely white pixel values at all positions) were discarded from the analysis. Each layer of the RGB image matrix was separated and converted to a greyscale image. Putative Mtb bacilli were then manually identified and used to set threshold values to eliminate background noise relative to Mtb signal. Thresholded images were then used to identify “Mtb-like” objects in an image for further downstream verification. Each of these objects was numerically labelled and compared against a user determined spectral profile to identify putative Mtb and eliminate false positives. Host cellular nuclei were identified in an identical fashion, but compared to a different spectral profile to identify true positives. Mtb and cellular nuclear objects identified in this way were tested for association, based on mean alveolar macrophage radius [47], and added to a matrix that was mapped onto a reduced image of the whole tissue section to reveal physical locations of Mtb infection. Following automated Mtb object detection, each object was manually validated to ensure only putative detections were used for downstream analysis. Additionally, aggregated or single Mtb classification was manually validated. The resultant curated matrices were exported to R for graphing. Average single bacterium size was calculated by finding the mean area of manually validated Mtb single bacteria. Mean single Mtb area was then used to estimate the number of bacteria in all Mtb objects. The largest manually validated single mtb object was used as a threshold above which all other Mtb objects were classified as aggregates.

### Macrophage cultures

Peripheral blood mononuclear cells were isolated by density gradient centrifugation using Histopaque 1077 (Sigma-Aldrich, St Louis, MO). CD14+ monocytes were purified under positive selection using anti-CD14 microbeads (Miltenyi Biotec, San Diego, CA). For RNA-Seq protocols, CD14+ monocytes were seeded at 1 *×* 10^6^ cells per well in non-tissue culture treated 35 mm 6-well plates. For time-lapse microscopy protocols, CD14+ monocytes were seeded at 0.2 *×* 10^6^ cells per 0.01% fibronectin (Sigma-Aldrich) coated 35 mm glass bottom optical dishes (Mattek, Ashland, MA). Monocytes were then differentiated in macrophage growth medium containing 1% each of HEPES, sodium pyruvate, L-glutamine, and non-essential amino acids, 10% human AB serum (Sigma-Aldrich), and 50 ng/ml GM-CSF (Peprotech, Rocky Hill, NJ) in RPMI. The cell culture medium was replaced one, three and six days post plating.

### Mtb Culture and macrophage infection

The mCherry fluorescent strain of H37Rv Mtb was derived by transforming the parental strain with a plasmid with mCherry under the smyc’ promoter (gift from D. Russell). Mtb were maintained in Difco Middlebrook 7H9 medium enriched with oleic acid-albumin-dextrose catalase supplement (BD, Sparks, MD). Three days before macrophage infection, Mtb were switched to grow in Tween 80-free media. On the day of infection, exponentially growing bacterial culture was pelleted at 2000 *×* g for 10 min, washed twice with 10 ml PBS, and large aggregates broken up by shaking with sterilized 2–4 mm glass beads for 30 s (bead beating). 10 ml of PBS was added and large clumps were further excluded by allowing them to settle for 5 min. Where heat killing was required, Mtb suspension was placed in a heating block for 20 minutes at 80 °C. To generate single Mtb bacilli, the bacterial suspension was passed through a 5 *µ*m syringe filter following aggregate preparation. The resulting singlet and aggregate Mtb suspensions were immediately used to infect differentiated MDM. Mtb grown in media containing Tween 80 surfactant was grown in parallel with detergent free Mtb culture to monitor bacterial growth and calibrate macrophage infection using optical density readings. MDM were infected with 150*µ*l Mtb aggregate suspension or 1000*µ*l singlet suspension for 3 hours, washed with PBS to remove extracellular Mtb and incubated for a further 3 hours.

### Cell sorting

Following infection, macrophages were lifted from non-tissue culture treated plates, using 1ml of Accutase (Sigma-Aldrich) cell dissociation reagent per 35mm well, and transferred to FACS tubes. 1*µ*l of Draq7 was added per macrophage containing FACS tube. Macrophage populations were gated into high and low Mtb infected populations based on Mtb mCherry fluorescence within each macrophage (measured at 561nm). Live and infected macrophages were selected for by gating out dead cells with compromised membranes using Draq7 fluorescence (at 633nm excitation). 10000 infected macrophages, per tube, were sorted into Trizol (Thermo-Fischer) and snap frozen in a dry ice and 99 percent isopropanol slurry using a BD FACSAria III flow cytometer (BD, New Jersey, US).

### RNA-Seq

Snap frozen samples were stored at −80C prior to transport and sequencing. Sample cDNA libraries were prepared according to protocols established by John J. Trombetta et.al [48] and sequenced on an Illumina NextSeq 500/550 instrument. Transcripts were aligned to human reference hg19 and read count libraries generated using the RSEM software package.

### Transcriptomics data analysis

Read count libraries, generated for each of the 15 replicates per treatment at an average read depth of 4 per base [49], were processed using the DESeq2 package for the R programming platform [50]. Metadata and read count matrices from each batch were concatenated into a single metadata and read count matrix prior to processing. For PCA analyses count matrices were R-log normalized in DESeq2, to prevent low abundance transcripts from dominating variation, and corrected for batch effects using the SVA ComBat function for R. Read count matrices were then arranged in descending order by variance across treatment conditions, and the top 0.1% of variable genes plotted in a PCA to reveal any clustering evident at the early assay timepoint. For differential expression analysis, the conventional DESeq2 negative binomial model was used, including blood donor as a factor. Candidate genes identified in this way were cross-referenced with the R-Log normalised, variance ordered lists to narrow potential candidate gene lists. Genes identified were then tested for significant differences between infection conditions using a Bonferronni corrected Mann-Whitney U-test.

### Cytokine analysis

MDMs were isolated and differentiated as previously described, at a concentration of 1 *×* 10^6^ cells per well on tissue culture treated 35 mm 6-well plates. MDMs were infected with 150*µ*l Mtb aggregate suspension or 1000*µ*l Mtb single suspension and incubated for 3h before being washed with PBS to remove extracellular Mtb, and incubated for a further 3h. Supernatant was then collected, 0.2*µ*m filtered and frozen prior to cytokine quantification. Cytokine levels were quantified using a Biorad Bioplex 200 (BioRad, California, US) instrument and custom R&D systems Luminex cytokine panel kit, according to kit instructions. Custom panels were constructed to have a broad array of cytokines to validate against transcriptional data. Due to the early supernatant harvesting timepoint, only cytokines that had levels above background were retained for analysis purposes.

### Microscopy

Macrophages and bacteria were imaged using an Andor (Andor, Belfast, UK) integrated Metamorph-controlled (Molecular Devices, Sunnyvale, CA) Nikon TiE motorized microscope (Nikon Corporation, Tokyo, Japan) with a 20x, 0.75 NA phase objective. Images were captured using an 888 EMCCD camera (Andor). Temperature and C0_2_ were maintained at 37C and 5% using an environmental chamber (OKO Labs, Naples, Italy).For timelapse protocols, images were captured once every ten minutes for the duration of the time-lapse. For each acquisition, images were captured at wavelengths applicable to fluorophores used in the analysis including transmitted light (phase contrast), 561nm (RFP), and 640nm (DRAQ7, lysotracker). Image analysis was performed using custom written matlab script. Single cell segmentation was manually carried out prior to fluorescent signal quantification. For each cell, fluorescent signal in each channel was quantified as pixel intensity.

### Macrophage acidification assay

Single cell fluorescence data for lysotracker acidification was acquired at a single timepoint at 6 hours post infection using the confocal microscopy system described previously. MDM on fibronectin coated optical dishes were infected with 400*µ*l Mtb aggregate suspension and incubated for 3 hours before being washed with PBS to remove any cell free Mtb and further incubated for 2 hours. 1 hour before image acquisition, Lysotracker was added to wells at a concentration of 75 nM. Images were processed as previously described to acquire pixel fluorescence intensity data for each fluorescent channel per cell. Model fit was to 3*/r*, where *r* was aggregate radius.

## Acknowledgments

This study was supported by a Bill and Melinda Gates Foundation Award OPP1116944. IMF was supported through a Sub-Saharan African Network for TB/HIV Research Excellence (SANTHE, a DELTAS Africa Initiative (grant DEL-15–006)) fellowship. AKS was supported by the Searle Scholars Program, the Beckman Young Investigator Program, and a Sloan Fellowship in Chemistry.

## Supplementary figures

**Figure S1:**
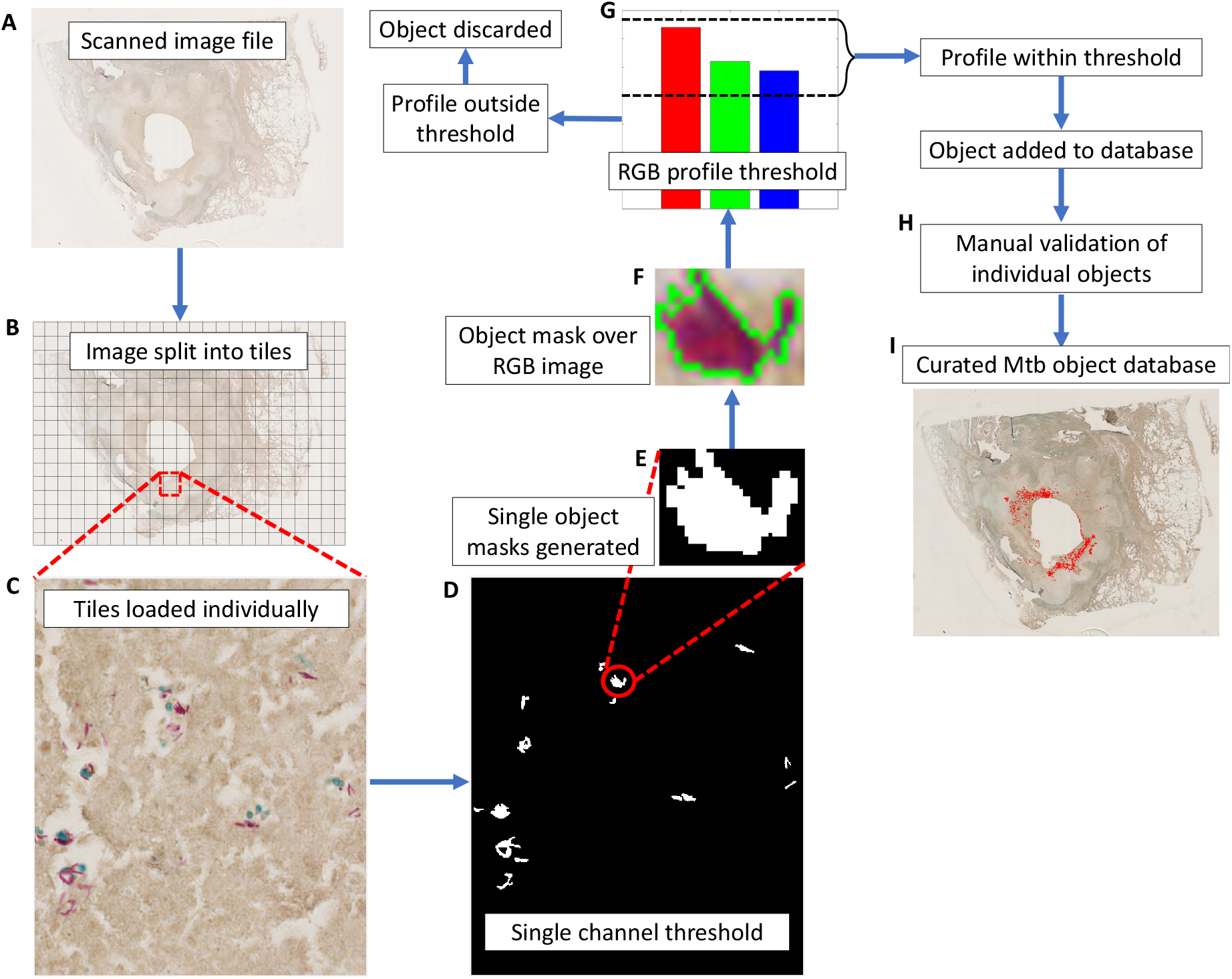
Schematic overview of semi-automated histological image analysis. (A) The section was scanned using a Hamamatsu Nanozoomer 2.0 rs slide scanner and exported to (B) ImageJ to split large files into smaller image tiles in preparation for processing in Matlab.(C) Smaller image tiles were individually loaded into Matlab and (D) thresholded in the Mtb channel to create binary masks corresponding to the locations of Mtb bacilli. Each of these individual binary object masks (E) were applied to the original RGB image (F) to isolate full RGB profiles of the objects. Each of these object profiles was then compared to a reference RGB pattern that matched the RGB profile of stained Mtb (G). Objects that were within the RGB thresholds were added to the database. Objects that succeeded the RGB profile thresholds were then individually manually curated (H) and added to the curated database (I).

**Table S1:** GSEA differentially regulated gene sets between infection conditions at Nominal p-value and FDR *<* 0.05.

| Gene set | Gene set size | NES | Nominal P-value | FDR q-value |
| --- | --- | --- | --- | --- |
| **Aggregate vs. Uninfected** |  |  |  |  |
| tnfa signaling via nfkb | 200 | 2,81 | <0,001 | <0,001 |
| inflammatory response | 200 | 2,46 | <0,001 | <0,001 |
| allograft rejection | 200 | 1,63 | <0,001 | 0,010 |
| interferon gamma response | 200 | 1,61 | <0,001 | 0,009 |
| il6 jak stat3 signaling | 87 | 1,61 | <0,001 | 0,007 |
| hypoxia | 200 | 1,59 | <0,001 | 0,008 |
| cholesterol homeostasis | 74 | 1,46 | 0,009 | 0,038 |
| apoptosis | 161 | 1,44 | <0,001 | 0,038 |
| kras signaling up | 200 | 1,43 | 0,004 | 0,037 |
| complement | 200 | 1,39 | 0,003 | 0,046 |
| **Multiple vs. Uninfected** |  |  |  |  |
| tnfa signaling via nfkb | 200 | 2,59 | <0,001 | <0,001 |
| inflammatory response | 200 | 2,24 | <0,001 | <0,001 |
| il6 jak stat3 signaling | 87 | 1,81 | <0,001 | <0,001 |
| complement | 200 | 1,59 | <0,001 | 0,014 |
| interferon gamma response | 200 | 1,58 | <0,001 | 0,013 |
| allograft rejection | 200 | 1,57 | <0,001 | 0,012 |
| apoptosis | 161 | 1,45 | 0,004 | 0,043 |
| **Single vs Uninfected** |  |  |  |  |
| tnfa signaling via nfkb | 200 | 2,42 | <0,001 | <0,001 |
| inflammatory response | 200 | 1,83 | <0,001 | <0,001 |
| cholesterol homeostasis | 74 | 1,57 | 0,011 | 0,032 |
| notch signaling | 32 | 1,51 | 0,025 | 0,038 |
| complement | 200 | 1,48 | <0,001 | 0,040 |
| **Multiple vs. Single** |  |  |  |  |
| inflammatory response | 200 | 1,73 | <0,001 | 0,004 |
| tnfa signaling via nfkb | 200 | 1,68 | <0,001 | 0,004 |
| il6 jak stat3 signaling | 87 | 1,63 | <0,001 | 0,005 |
| e2f targets | 200 | 1,46 | 0,001 | 0,040 |
| **Aggregate vs. Single** |  |  |  |  |
| tnfa signaling via nfkb | 200 | 2,24 | <0,001 | <0,001 |
| inflammatory response | 200 | 1,87 | <0,001 | 0,001 |
| **Aggregate vs. Multiple** |  |  |  |  |
| tnfa signaling via nfkb | 200 | 2,07 | <0,001 | <0,001 |

**Table S2:**
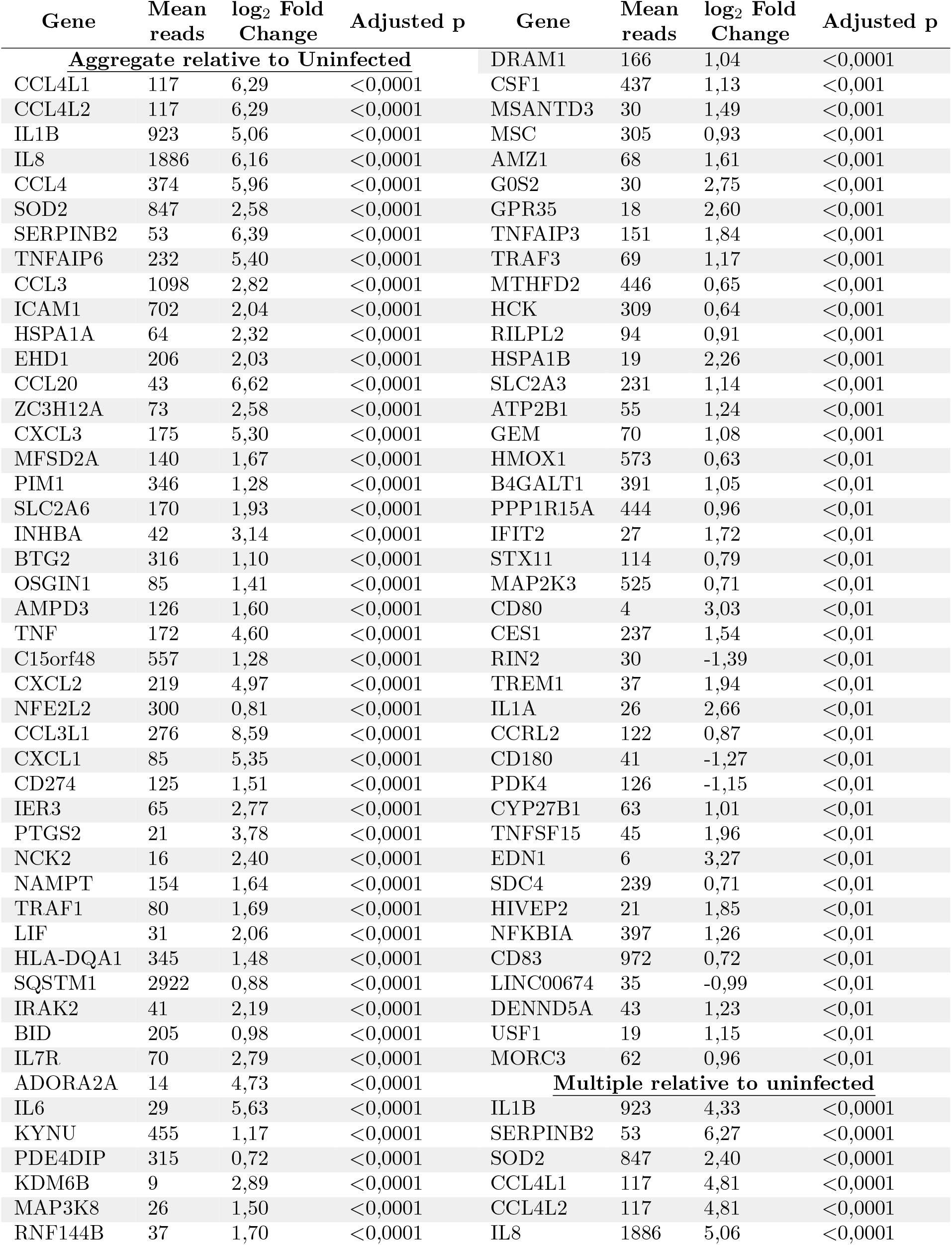
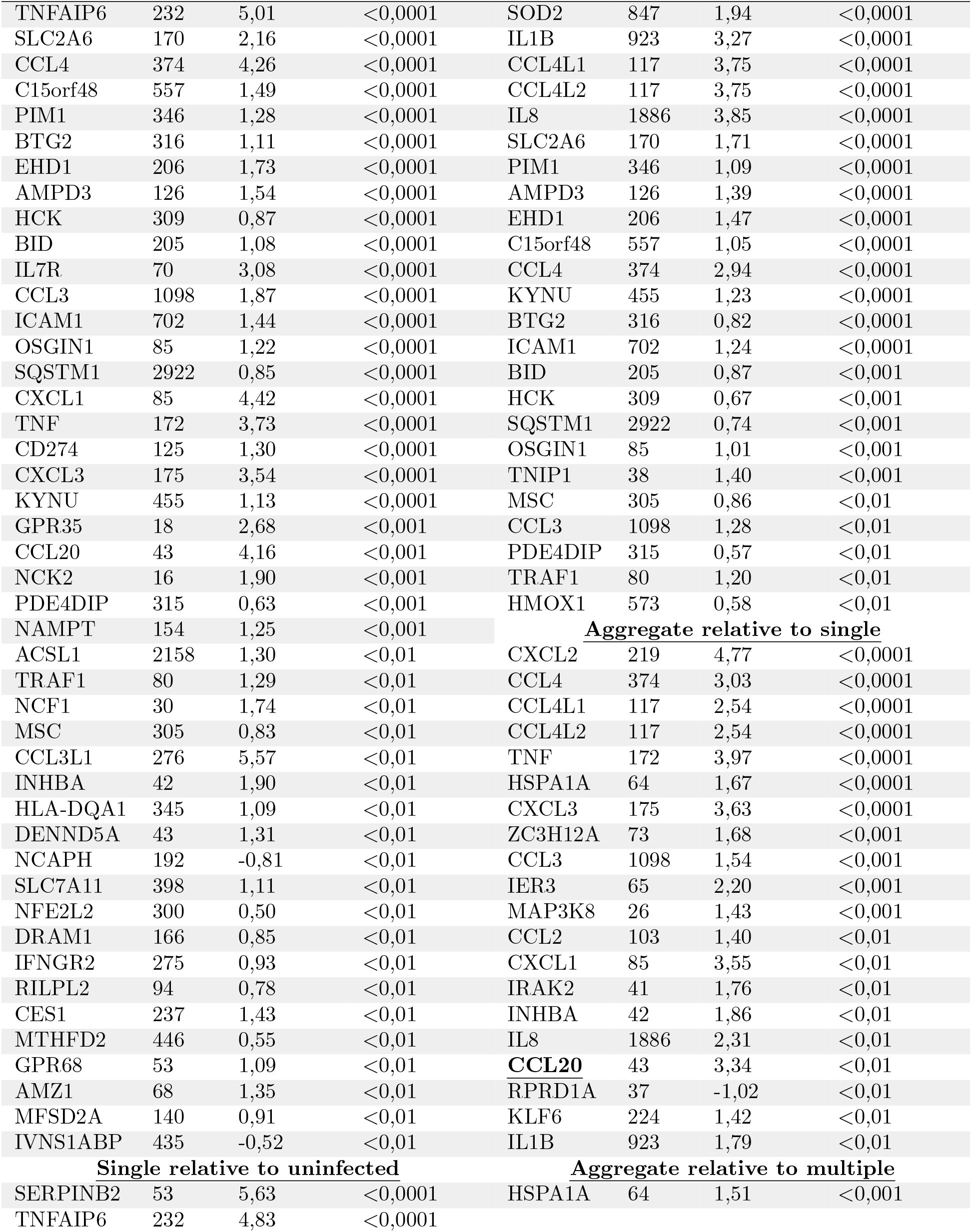
DESeq2 differentially regulated genes between infection conditions at adjusted p-value *<* 0.01

| Gene | Mean reads | log<sub>2</sub> Fold Change | Adjusted p | Gene | Mean reads | log<sub>2</sub> Fold Change | Adjusted p |
| --- | --- | --- | --- | --- | --- | --- | --- |
| **Aggregate relative to Uninfected** |  |  |  | DRAM1 | 166 | 1,04 | <0,0001 |
| CCL4L1 | 117 | 6,29 | <0,0001 | CSF1 | 437 | 1,13 | <0,001 |
| CCL4L2 | 117 | 6,29 | <0,0001 | MSANTD3 | 30 | 1,49 | <0,001 |
| IL1B | 923 | 5,06 | <0,0001 | MSC | 305 | 0,93 | <0,001 |
| IL8 | 1886 | 6,16 | <0,0001 | AMZ1 | 68 | 1,61 | <0,001 |
| CCL4 | 374 | 5,96 | <0,0001 | G0S2 | 30 | 2,75 | <0,001 |
| SOD2 | 847 | 2,58 | <0,0001 | GPR35 | 18 | 2,60 | <0,001 |
| SERPINB2 | 53 | 6,39 | <0,0001 | TNFAIP3 | 151 | 1,84 | <0,001 |
| TNFAIP6 | 232 | 5,40 | <0,0001 | TRAF3 | 69 | 1,17 | <0,001 |
| CCL3 | 1098 | 2,82 | <0,0001 | MTHFD2 | 446 | 0,65 | <0,001 |
| ICAM1 | 702 | 2,04 | <0,0001 | HCK | 309 | 0,64 | <0,001 |
| HSPA1A | 64 | 2,32 | <0,0001 | RILPL2 | 94 | 0,91 | <0,001 |
| EHD1 | 206 | 2,03 | <0,0001 | HSPA1B | 19 | 2,26 | <0,001 |
| CCL20 | 43 | 6,62 | <0,0001 | SLC2A3 | 231 | 1,14 | <0,001 |
| ZC3H12A | 73 | 2,58 | <0,0001 | ATP2B1 | 55 | 1,24 | <0,001 |
| CXCL3 | 175 | 5,30 | <0,0001 | GEM | 70 | 1,08 | <0,001 |
| MFSD2A | 140 | 1,67 | <0,0001 | HMOX1 | 573 | 0,63 | <0,01 |
| PIM1 | 346 | 1,28 | <0,0001 | B4GALT1 | 391 | 1,05 | <0,01 |
| SLC2A6 | 170 | 1,93 | <0,0001 | PPP1R15A | 444 | 0,96 | <0,01 |
| INHBA | 42 | 3,14 | <0,0001 | IFIT2 | 27 | 1,72 | <0,01 |
| BTG2 | 316 | 1,10 | <0,0001 | STX11 | 114 | 0,79 | <0,01 |
| OSGIN1 | 85 | 1,41 | <0,0001 | MAP2K3 | 525 | 0,71 | <0,01 |
| AMPD3 | 126 | 1,60 | <0,0001 | CD80 | 4 | 3,03 | <0,01 |
| TNF | 172 | 4,60 | <0,0001 | CES1 | 237 | 1,54 | <0,01 |
| C15orf48 | 557 | 1,28 | <0,0001 | RIN2 | 30 | -1,39 | <0,01 |
| CXCL2 | 219 | 4,97 | <0,0001 | TREM1 | 37 | 1,94 | <0,01 |
| NFE2L2 | 300 | 0,81 | <0,0001 | IL1A | 26 | 2,66 | <0,01 |
| CCL3L1 | 276 | 8,59 | <0,0001 | CCRL2 | 122 | 0,87 | <0,01 |
| CXCL1 | 85 | 5,35 | <0,0001 | CD180 | 41 | -1,27 | <0,01 |
| CD274 | 125 | 1,51 | <0,0001 | PDK4 | 126 | -1,15 | <0,01 |
| IER3 | 65 | 2,77 | <0,0001 | CYP27B1 | 63 | 1,01 | <0,01 |
| PTGS2 | 21 | 3,78 | <0,0001 | TNFSF15 | 45 | 1,96 | <0,01 |
| NCK2 | 16 | 2,40 | <0,0001 | EDN1 | 6 | 3,27 | <0,01 |
| NAMPT | 154 | 1,64 | <0,0001 | SDC4 | 239 | 0,71 | <0,01 |
| TRAF1 | 80 | 1,69 | <0,0001 | HIVEP2 | 21 | 1,85 | <0,01 |
| LIF | 31 | 2,06 | <0,0001 | NFKBIA | 397 | 1,26 | <0,01 |
| HLA-DQA1 | 345 | 1,48 | <0,0001 | CD83 | 972 | 0,72 | <0,01 |
| SQSTM1 | 2922 | 0,88 | <0,0001 | LINC00674 | 35 | -0,99 | <0,01 |
| IRAK2 | 41 | 2,19 | <0,0001 | DENND5A | 43 | 1,23 | <0,01 |
| BID | 205 | 0,98 | <0,0001 | USF1 | 19 | 1,15 | <0,01 |
| IL7R | 70 | 2,79 | <0,0001 | MORC3 | 62 | 0,96 | <0,01 |
| ADORA2A | 14 | 4,73 | <0,0001 | **Multiple relative to uninfected** |  |  |  |
| IL6 | 29 | 5,63 | <0,0001 | IL1B | 923 | 4,33 | <0,0001 |
| KYNU | 455 | 1,17 | <0,0001 | SERPINB2 | 53 | 6,27 | <0,0001 |
| PDE4DIP | 315 | 0,72 | <0,0001 | SOD2 | 847 | 2,40 | <0,0001 |
| KDM6B | 9 | 2,89 | <0,0001 | CCL4L1 | 117 | 4,81 | <0,0001 |
| MAP3K8 | 26 | 1,50 | <0,0001 | CCL4L2 | 117 | 4,81 | <0,0001 |
| RNF144B | 37 | 1,70 | <0,0001 | IL8 | 1886 | 5,06 | <0,0001 |

| Gene | Mean reads | log2 Fold Change | Adjusted p | Gene | Mean reads | log2 Fold Change | Adjusted p |
| --- | --- | --- | --- | --- | --- | --- | --- |
| TNFAIP6 | 232 | 5,01 | <0,0001 | SOD2 | 847 | 1,94 | <0,0001 |
| SLC2A6 | 170 | 2,16 | <0,0001 | IL1B | 923 | 3,27 | <0,0001 |
| CCL4 | 374 | 4,26 | <0,0001 | CCL4L1 | 117 | 3,75 | <0,0001 |
| C15orf48 | 557 | 1,49 | <0,0001 | CCL4L2 | 117 | 3,75 | <0,0001 |
| PIM1 | 346 | 1,28 | <0,0001 | IL8 | 1886 | 3,85 | <0,0001 |
| BTG2 | 316 | 1,11 | <0,0001 | SLC2A6 | 170 | 1,71 | <0,0001 |
| EHD1 | 206 | 1,73 | <0,0001 | PIM1 | 346 | 1,09 | <0,0001 |
| AMPD3 | 126 | 1,54 | <0,0001 | AMPD3 | 126 | 1,39 | <0,0001 |
| HCK | 309 | 0,87 | <0,0001 | EHD1 | 206 | 1,47 | <0,0001 |
| BID | 205 | 1,08 | <0,0001 | C15orf48 | 557 | 1,05 | <0,0001 |
| IL7R | 70 | 3,08 | <0,0001 | CCL4 | 374 | 2,94 | <0,0001 |
| CCL3 | 1098 | 1,87 | <0,0001 | KYNU | 455 | 1,23 | <0,0001 |
| ICAM1 | 702 | 1,44 | <0,0001 | BTG2 | 316 | 0,82 | <0,0001 |
| OSGIN1 | 85 | 1,22 | <0,0001 | ICAM1 | 702 | 1,24 | <0,0001 |
| SQSTM1 | 2922 | 0,85 | <0,0001 | BID | 205 | 0,87 | <0,001 |
| CXCL1 | 85 | 4,42 | <0,0001 | HCK | 309 | 0,67 | <0,001 |
| TNF | 172 | 3,73 | <0,0001 | SQSTM1 | 2922 | 0,74 | <0,001 |
| CD274 | 125 | 1,30 | <0,0001 | OSGIN1 | 85 | 1,01 | <0,001 |
| CXCL3 | 175 | 3,54 | <0,0001 | TNIP1 | 38 | 1,40 | <0,001 |
| KYNU | 455 | 1,13 | <0,0001 | MSC | 305 | 0,86 | <0,01 |
| GPR35 | 18 | 2,68 | <0,001 | CCL3 | 1098 | 1,28 | <0,01 |
| CCL20 | 43 | 4,16 | <0,001 | PDE4DIP | 315 | 0,57 | <0,01 |
| NCK2 | 16 | 1,90 | <0,001 | TRAF1 | 80 | 1,20 | <0,01 |
| PDE4DIP | 315 | 0,63 | <0,001 | HMOX1 | 573 | 0,58 | <0,01 |
| NAMPT | 154 | 1,25 | <0,001 | **Aggregate relative to single** |  |  |  |
| ACSL1 | 2158 | 1,30 | <0,01 | CXCL2 | 219 | 4,77 | <0,0001 |
| TRAF1 | 80 | 1,29 | <0,01 | CCL4 | 374 | 3,03 | <0,0001 |
| NCF1 | 30 | 1,74 | <0,01 | CCL4L1 | 117 | 2,54 | <0,0001 |
| MSC | 305 | 0,83 | <0,01 | CCL4L2 | 117 | 2,54 | <0,0001 |
| CCL3L1 | 276 | 5,57 | <0,01 | TNF | 172 | 3,97 | <0,0001 |
| INHBA | 42 | 1,90 | <0,01 | HSPA1A | 64 | 1,67 | <0,0001 |
| HLA-DQA1 | 345 | 1,09 | <0,01 | CXCL3 | 175 | 3,63 | <0,0001 |
| DENND5A | 43 | 1,31 | <0,01 | ZC3H12A | 73 | 1,68 | <0,001 |
| NCAPH | 192 | -0,81 | <0,01 | CCL3 | 1098 | 1,54 | <0,001 |
| SLC7A11 | 398 | 1,11 | <0,01 | IER3 | 65 | 2,20 | <0,001 |
| NFE2L2 | 300 | 0,50 | <0,01 | MAP3K8 | 26 | 1,43 | <0,001 |
| DRAM1 | 166 | 0,85 | <0,01 | CCL2 | 103 | 1,40 | <0,01 |
| IFNGR2 | 275 | 0,93 | <0,01 | CXCL1 | 85 | 3,55 | <0,01 |
| RILPL2 | 94 | 0,78 | <0,01 | IRAK2 | 41 | 1,76 | <0,01 |
| CES1 | 237 | 1,43 | <0,01 | INHBA | 42 | 1,86 | <0,01 |
| MTHFD2 | 446 | 0,55 | <0,01 | IL8 | 1886 | 2,31 | <0,01 |
| GPR68 | 53 | 1,09 | <0,01 | **CCL20** | 43 | 3,34 | <0,01 |
| AMZ1 | 68 | 1,35 | <0,01 | RPRD1A | 37 | -1,02 | <0,01 |
| MFSD2A | 140 | 0,91 | <0,01 | KLF6 | 224 | 1,42 | <0,01 |
| IVNS1ABP | 435 | -0,52 | <0,01 | IL1B | 923 | 1,79 | <0,01 |
| **Single relative to uninfected** |  |  |  | **Aggregate relative to multiple** |  |  |  |
| SERPINB2 | 53 | 5,63 | <0,0001 | HSPA1A | 64 | 1,51 | <0,001 |
| TNFAIP6 | 232 | 4,83 | <0,0001 |  |  |  |  |

